# Identification of high confidence human poly(A) RNA isoform scaffolds using nanopore sequencing

**DOI:** 10.1101/2020.11.18.389049

**Authors:** Logan Mulroney, Madalee G. Wulf, Ira Schildkraut, George Tzertzinis, John Buswell, Miten Jain, Hugh Olsen, Mark Diekhans, Ivan R. Corrêa, Mark Akeson, Laurence Ettwiller

**Affiliations:** Biomolecular Engineering Department, UC Santa Cruz, CA 95064; New England Biolabs, Ipswich, MA 01938; Genomics Institute, UC Santa Cruz, CA 95064

**Author notes:** First authors (Logan Mulroney, Madalee G. Wulf).

**Keywords:** RNA, transcriptomics, nanopore, Direct RNA, single-molecule, full-length, transcription start site, RNA isoforms

## Abstract

Nanopore sequencing devices read individual RNA strands directly. This facilitates identification of exon linkages and nucleotide modifications; however, using conventional methods the 5′ and 3′ ends of poly(A) RNA cannot be identified unambiguously. This is due in part to the architecture of the nanopore/enzyme-motor complex, and in part to RNA degradation *in vivo* and *in vitro* that can obscure transcription start and end sites. In this study, we aimed to identify individual full-length human RNA isoform scaffolds among ∼4 million nanopore poly(A)-selected RNA reads. First, to identify RNA strands bearing 5′ m^7^G caps, we exchanged the biological cap for a modified cap attached to a 45-nucleotide oligomer. This oligomer adaptation method improved 5′ end sequencing and ensured correct identification of the 5′ m^7^G capped ends. Second, among these 5′-capped nanopore reads, we screened for ionic current signatures consistent with a 3′ polyadenylation site. Combining these two steps, we identified 294,107 individual high-confidence full-length RNA scaffolds, most of which (257,721) aligned to protein-coding genes. Of these, 4,876 scaffolds indicated unannotated isoforms that were often internal to longer, previously identified RNA isoforms. Orthogonal data confirmed the validity of these high-confidence RNA scaffolds.

## INTRODUCTION

Most human genes encode multiple transcript isoforms. These isoforms are derived from alternative splicing, alternative transcription start sites (TSS), or alternative transcription termination sites (TTS). TSS outnumber genes^1^, and together with alternate TTS account for most transcript isoform differences between tissues^2^. Accurate identification of an RNA isoform is difficult when either its TSS or its TTS is unknown or positioned within the genomic region of a larger isoform^3^, and internal isoforms are often omitted from transcriptome annotations^4^. Direct sequencing of nucleotides between the 5′ cap and 3′ poly(A) tail on individual RNA molecules would reveal the isoform structure and associated modifications without employing error-prone computational tools^5–10^.

Nanopore RNA sequencing is a single molecule technique that reads RNA directly rather than cDNA copies^11–13^. This avoids cDNA artifacts^14^ and permits detection of RNA modifications, thus far including m^6^A^11–13^, inosine^13^, pseudouridine^15^, and m^7^G^15^. Approximate poly(A) tail lengths can also be discerned for those reads^13^.

However, using standard protocols, nanopore direct RNA reads terminate before reaching the 5′ end of captured molecules. This is because the enzyme that regulates translocation prematurely releases captured strands^13^. Accurate identification of full-length isoforms is further complicated by RNA strand degradation that occurs in the cell, during sample preparation, or in silico^13^. A possible marker for full-length reads would be the 5′ m^7^G cap that is associated with eukaryotic mRNA^16^. Two studies documented the presence of the cap in nanopore reads^12,17^ using enzymatic decapping and ligation^18^.

Previously, we successfully replaced the m^7^G caps with biotin-modified caps to enrich for native 5′ mRNA ends which enabled the identification of high confidence TSS using Illumina sequencing^19^. In this study, we introduce a parallel chemo-enzymatic method, Nanopore ReCappable sequencing (NRCeq), that uniquely and specifically replaces m^7^G caps with an RNA oligonucleotide adapter. This approach avoids common shortfalls of ligation methods.

We used NRCeq to identify individual full-length high-confidence RNA scaffolds in a GM12878 poly(A) RNA transcriptome. These scaffolds extended from the m^7^G cap to a documented poly(A) site. We systematically correlated these scaffolds with orthogonal data and showed that single reads could provide compelling evidence for *bona fide* RNA isoforms.

## RESULTS

### NRCeq strategy

The NRCeq strategy is diagrammed in Fig. 1a. First, we used the yeast scavenger decapping enzyme (yDcpS)^20,21^ to remove the m^7^G cap from poly(A)-enriched RNA, leaving 5′-diphosphate ends. Second, the 5′-diphosphate RNA strands were recapped with 3′-azido-ddGTP using *Vaccinia* capping enzyme^21^ (Supplementary Fig. 2). Third, the 3′-azido recapped RNA strands were covalently attached to a dibenzocyclooctyne (DBCO) reactive group on the 3′ end of a 45 nt long RNA oligonucleotide adapter using specific copper-free ‘click’ chemistry^22,23^ (Supplementary Fig.3).

**Figure 1.**
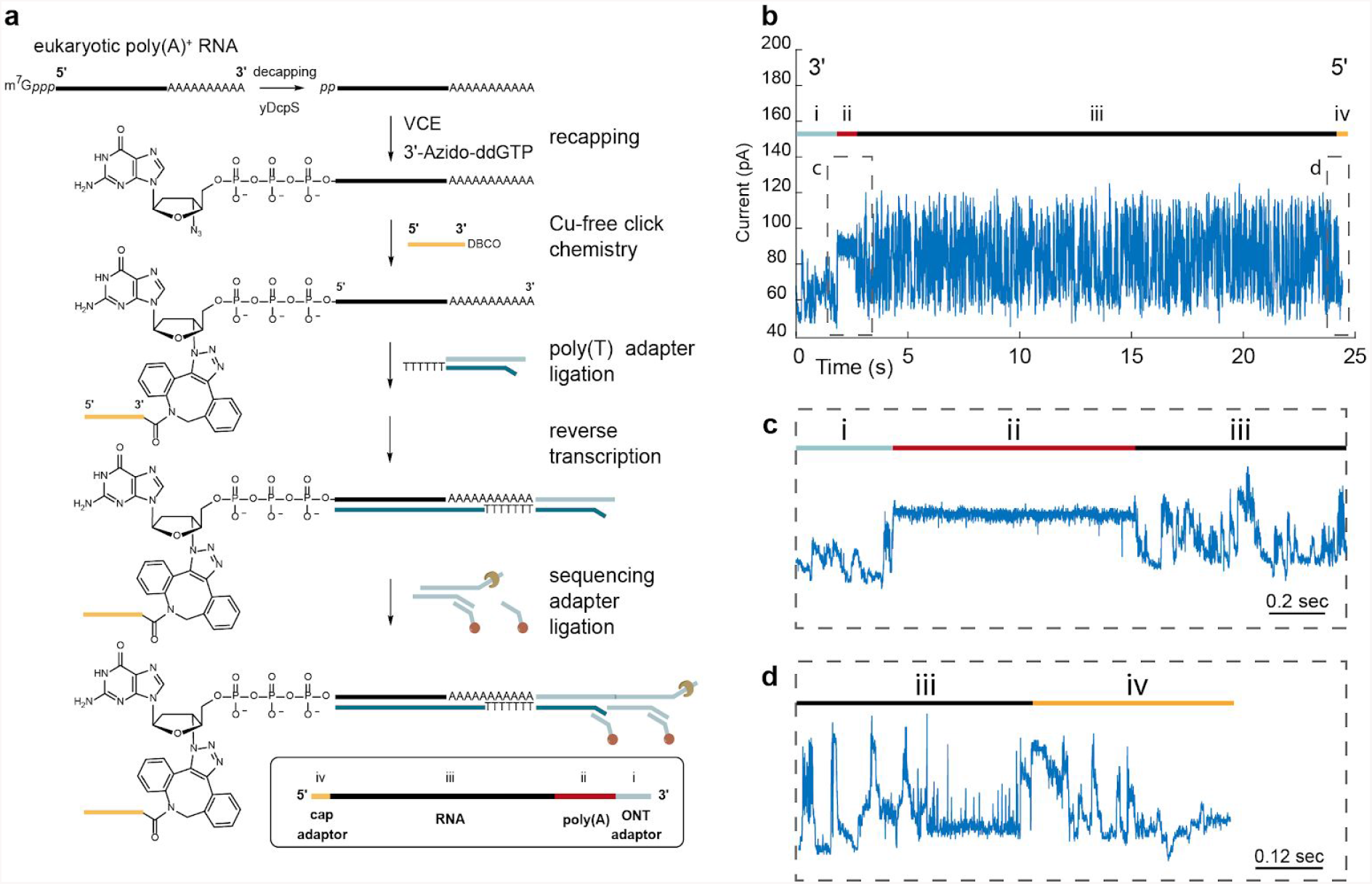
Preparation and analysis of 5′ cap-adapted poly(A) strands. (**a**) Adaptation and library preparation workflow for poly(A)-selected RNA. (**b**) Representative ionic current trace for a cap-adapted full-length RNA read is shown for the thymidine phosphorylase gene (TYMP). The trace begins with ionic current associated with the ONT adapter (i). This is followed by a monotonic ionic current associated with the 3′ poly(A) tail (ii) and then a variable ionic current associated with the RNA transcript nucleotides (iii). The final segment is an ionic current signature characteristic of the 45 nt RNA cap-adapter (iv). (**c**) An approximately two second window centered on the ionic current associated with the poly(A) tail (ii). (**d**) An approximately one second window centered on the boundary between the ionic current associated with the 5′ end of the transcript (iii) and a characteristic adapter ionic current trace (iv).

A typical cap-adapted ionic current trace for human thymidine phosphorylase (TYMP) is shown in Fig. 1b. Following strand capture, a characteristic ionic current is caused by translocation of the ONT 3′ adapter (i). This is followed by a monotonic ionic current associated with the 3′ poly(A) tail (ii) and then a variable ionic current with a bottle brush appearance associated with a mixed series of RNA nucleotides (iii). The trace terminated with an ionic current signature characteristic of the 45 nt RNA cap-adapter (iv). This signature indicated that individual strands were read from the 3′ poly(A) tail (Fig. 1c) through the original 5′-capped end (Fig. 1d). We used a sequence-based barcode identification software (Porechop https://github.com/rrwick/Porechop) to detect the adapter on individual nanopore reads (see Methods).

### Optimizing 5′ cap-adaptation using *Saccharomyces cerevisiae* poly(A) RNA

We optimized the 5′ cap-adaptation strategy using *S. cerevisiae* poly(A) RNA. The yeast transcriptome is well-suited for this because the m^7^G cap is identical to the human m^7^G cap, and because most yeast genes encode only one RNA isoform^24^.

Initially, we used a copper-catalyzed click reaction for the 5′ adaptation step (Supplementary Methods), however RNA degradation was unacceptable as measured by RNA integrity number (RIN)^25^ (Supplementary Table 2). As an alternative, we implemented a copper-free chemistry step based on a strain-promoted click reaction (Fig. 1a, and Methods)^23,26^. This eliminated RNA degradation during the click step (Supplementary Table 2). Further, this change from copper-catalyzed to copper-free chemistry improved the percentage of yeast poly(A) RNA reads that were cap-adapted (13.4% to 38.4% respectively), and the average strand read length (N50 equal to 692 to 744 nt respectively).

### Applying NRCeq to human poly(A) RNA transcripts

Having optimized 5′ cap-adaption chemistry and detection, we applied the cap-adaptation strategy to poly(A) RNA isolated from GM12878 cells, a model human B-lymphocyte cell line. We acquired 4 million reads with quality scores greater than or equal to Q7 that went through the cap-adaptation process (we refer to this population as ‘treated reads’ in the text that follows, see Fig. 6a). We identified 574,091 (14.3%) of the treated reads as ‘cap-adapted’ (see Methods, Fig 6b, and Supplementary Figure 1). As a control, we also performed standard native RNA nanopore sequencing using the same starting poly(A) RNA material (‘untreated reads’) yielding approximately 3.8 million reads from 2 replicates (Supplementary Table 1).

The N50 value for the cap-adapted reads was 1301 nt, which was shorter than the N50 value for untreated reads (1614 nt) (Supplementary Table 1). Given this difference, we were concerned that the cap-adaptation process adversely affected human RNA transcript recovery. To test this, we compared the number of transcript copies per gene for the untreated and treated samples. The Spearman rank correlation score was very strong (0.95), indicating that the treatment protocol did not substantially affect RNA strand recovery (Fig. 2a). We then compared the number of transcript copies per gene for the cap-adapted and full-length untreated samples using a previously described definition for full-length^13,15^. The Spearman rank correlation score was lower, but also very strong (0.83) (Fig. 2b).

**Figure 2.**
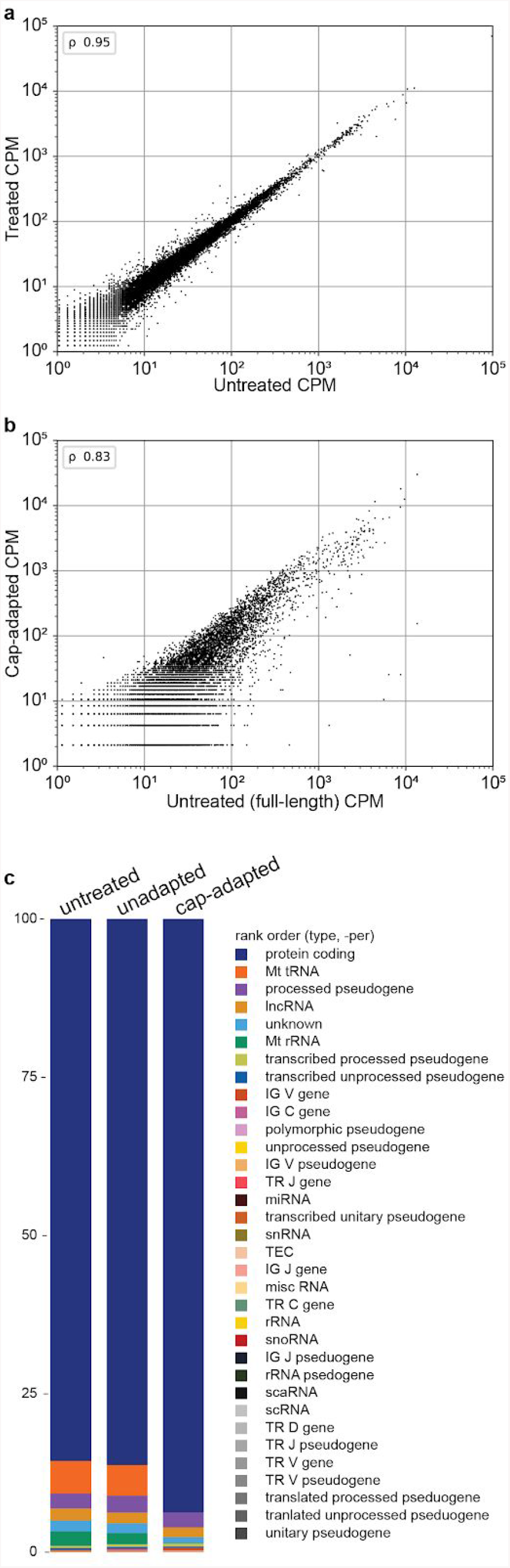
Cap-adaptation performance on GM12878 RNA. Number of transcripts per gene counts per million (CPM) correlation plots (**a**) between untreated and treated samples, and (**b**) between full-length untreated and cap-adapted samples. Pearson’s r (ρ) were 0.95 and 0.83, respectively. (**c**) Percent of RNA by class for untreated, unadapted, and cap-adapted reads. Unadapted refers to reads within the treated samples that are missing the adapter sequence. All class percentages are in Supplementary Table 3.

Among untreated and cap-adapted reads, 85% and 94% of aligned nanopore reads, respectively, correspond to protein-coding genes (Fig. 2c). Mitochondrial RNA reads accounted for 11% of the mapped untreated reads. By comparison, mitochondrial RNA reads account for only 0.3% of the mapped cap-adapted reads. This result was expected because mitochondrial transcript 5′ ends usually bear a 5′ monophosphate or are transient, with a subset capped by NAD^+^ and NADH^27^.

### TSS identification with NRCeq

NRCeq was designed to identify m^7^G-capped RNA 5′ ends and improve base calling near those ends. We predicted that cap-adapted nanopore reads would be enriched for 5′ ends proximal to TSS annotated by GENCODE^4^. This prediction was substantiated (Fig. 3a). We found that 99% of 5′ ends for cap-adapted reads were within a window of 300 bp of an annotated TSS, compared to 77% of the untreated reads, and 65% of the treated reads (Fig. 3a).

**Figure 3.**
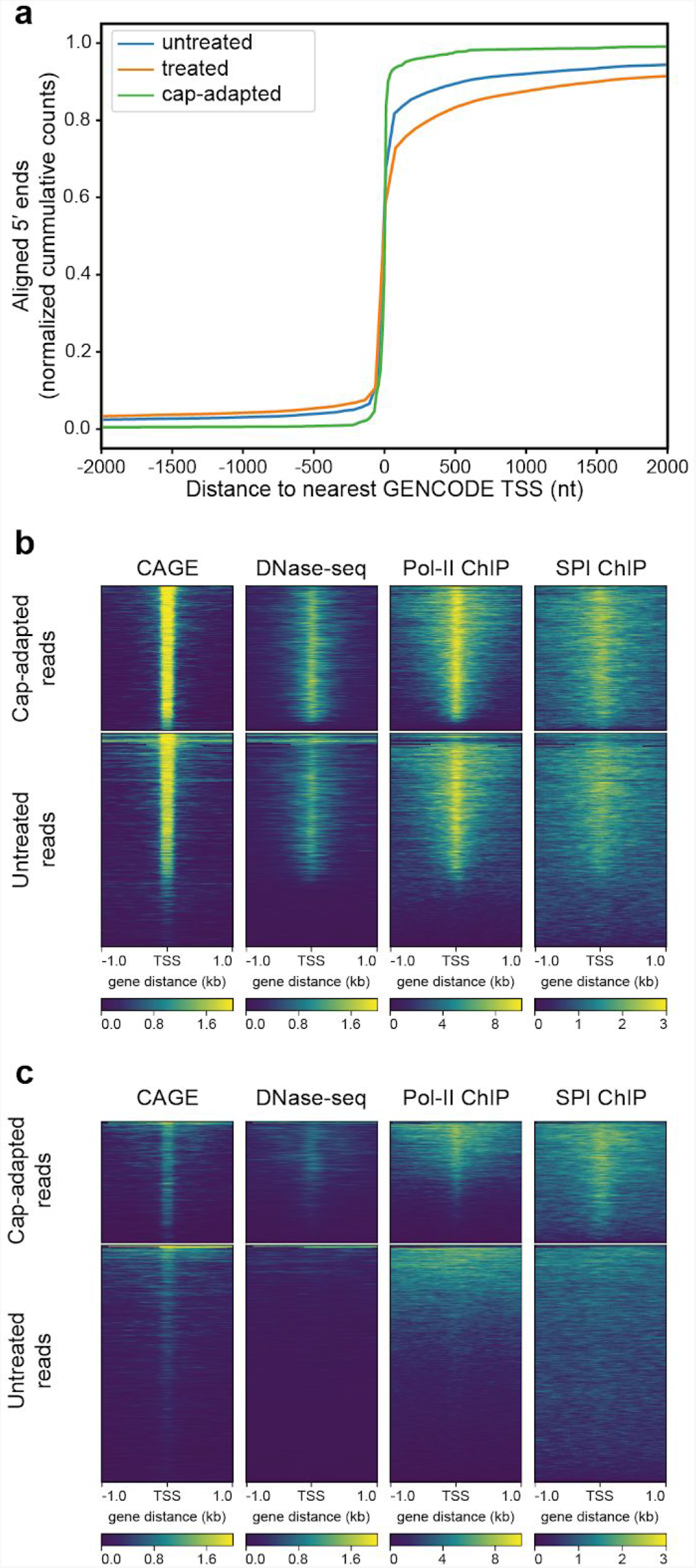
Correspondence between RNA nanopore 5′ end reads and orthogonal TSS evidence. (**a**) Nucleotide distance of RNA 5′ ends from annotated GENCODE TSS. The *x* axis is the number of nucleotides between a nanopore read 5′ end and the closest TSS annotated in GENCODE v.32. Negative numbers are upstream (5′) from the TSS; positive numbers are downstream (3′) from the TSS. The *y* axis is the cumulative number of RNA nanopore reads at a given distance of their 5′ end from an annotated GENCODE TSS. Values are normalized as a fraction of total counts for a given treatment. (**b**) Comparison of poly(A) RNA nanopore 5′ end reads to orthogonal TSS markers. Each plot is a heatmap where the *x* axis is a ±1 kb window centered on the 5′ end of each RNA nanopore read. Each row in the *y* axis is an individual read. The color intensity is the read depth normalized signal (CAGE, DNase-seq) or fold change over control for each position (POLR2 and SPI1). The top plots are cap-adapted reads, the bottom plots are untreated reads. (**c**) Comparison of unannotated RNA nanopore 5′ end reads to orthogonal TSS markers. Unannotated 5′ ends are defined as reads where the 5′ end is aligned more than 300 nucleotides away from an annotated TSS. The number of reads in each plot were downsampled to 9,116 reads (total number of unannotated cap-adapted reads).

Precise definition of TSS can be difficult. Therefore, we compared the start of the cap-adapted reads to those of other markers that are conventionally used for TSS determination. These included DNase-seq, Pol II Chip-seq, SPI1 ChiP-seq, and CAGE, all performed on the same cell line. We found that a majority of the cap-adapted reads corresponded with these other markers of transcription initiation (Fig. 3b).

It is noteworthy that a low number of the cap-adapted reads had 5′ ends that did not clearly map to annotated TSS, suggesting alternative TSS. To test this hypothesis, we filtered cap-adapted reads whose 5′ ends mapped >300 nt from any annotated GENCODE TSS^4^. We found 9,116 reads (1% of total) corresponding to 1,914 genes. A majority of these newly found 5′ ends were validated by DNase-seq, ChiP-seq, and CAGE (Fig. 3b), thus increasing the confidence that these are *bona fide* TSS^28^. The same pipeline identified 240,211 untreated reads (20% of total) that corresponded to unannotated 5′ TSS. However, the vast majority of these 5′ TSS were not validated by DNase-seq and ChiP-seq (Fig. 3c). Interestingly, a weak CAGE signal can be observed despite the lack of genomic TSS marks, which could be attributed to noise in the CAGE dataset.

### 5′ RACE validation of unannotated TSS

We used 5′ RACE^29^ to test the validity of 93 selected isoforms from 88 genes bearing presumptive new TSS (Fig. 4A). These TSS were chosen because they started either at an internal exon or at an unannotated exon. To eliminate processed monophosphate ends, total RNA was treated with CIP. To identify the position of capped ends, the RNA was also treated with RppH before library preparation to enable the enzymatic ligation of RNA oligonucleotide adapters to originally capped transcripts. Finally, to confirm the occurrence of the cap, we demonstrated the dependence of 5′ RACE products on decapping with RppH.

**Figure 4.**
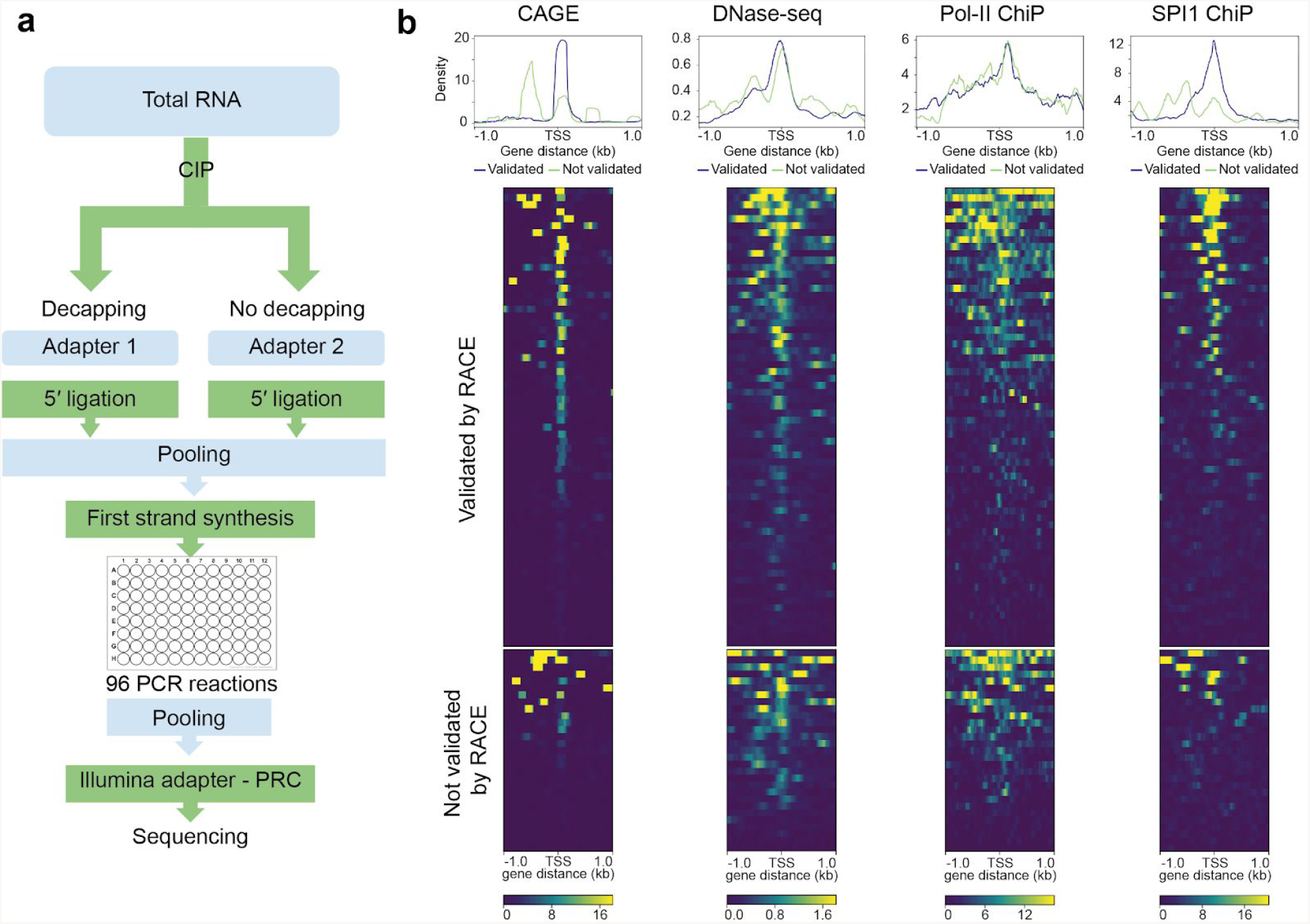
Multiplex RACE. **a**, Experimental design for RACE validation of 93 isoforms from 88 genes. **b**, Heatmaps of 3 promoter marks : PolR2 (ENCFF340BYJ) SPI1 (ENCFF289XSX) ChIP-seq, and DNase-seq (ENCFF093VXI) and CAGE (ENCFF580WIH), each showing 2 kb flanking regions at the 5′ end of the 93 candidate isoforms. The upper panel shows the 64 candidate TSS that were validated by RACE, the lower panel corresponds to the 29 TSS not validated by RACE.

Amplification involved a reverse PCR primer 100-200 nucleotides downstream of the predicted TSS. Our controls were two highly expressed genes with documented TSS, ACTB and TMSB10. Short-read 5′ RACE sequencing data confirmed 64 of the 93 target isoforms with RNA 5′ ends within 50 nucleotides of the cap-adapted TSS (Supplemental Table 4). Irrespective of the RACE results, CAGE or TSS chromatin marks corresponded with the 5′ end of many candidate scaffolds (Fig. 4B).

### Documentation of full-length RNA scaffolds

A central aim of this study was to facilitate mRNA isoform identification using individual full-length reads as scaffolds. This required identification of nanopore reads that aligned to protein-coding genes, and that correctly identified both the 5′ and 3′ ends of mature mRNA.

The 294,107 nanopore 5′ capped RNA reads identified by NRCeq were screened for the presence of poly(A) tails using nanopolish-polya^13^, which resulted in 222,687 reads. To filter for full-length RNA, we documented reads that had 3′ ends that aligned within -60 to +10 nt of annotated polyadenylation sites^30^ (Fig. 6e). This resulted in 209,093 individual full-length poly(A) RNA scaffolds (Fig. 5). There were 195,602 scaffolds that corresponded to 8,740 protein-coding genes (Fig. 6f). Per protein-coding gene transcript coverage ranged from 1-to-7,987 (https://github.com/mitenjain/dRNA_capping_analysis/blob/master/cap-adapted_full_length_scaffolds_geneCounts.txt). We identified 4,876 full-length RNA scaffolds with unannotated TSS that mapped to protein-coding genes (Fig 6g).

**Figure 5.**
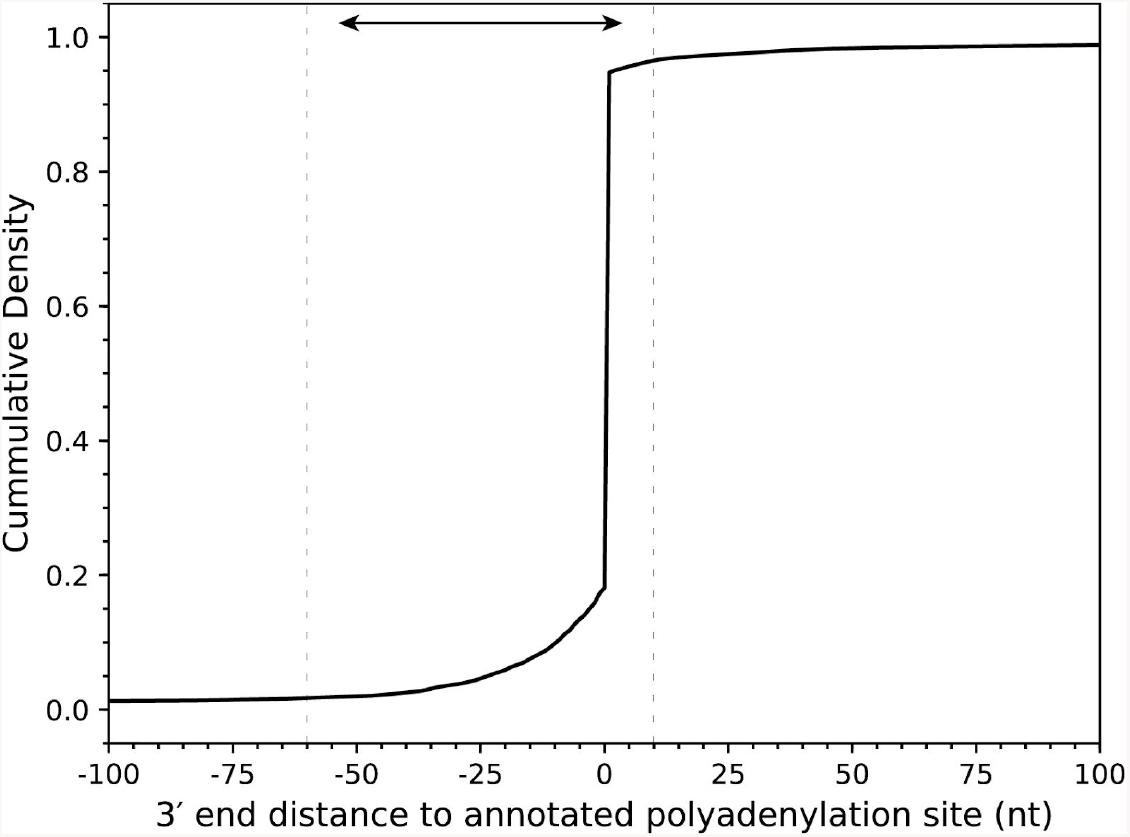
mRNA scaffold 3′ end distance to known polyadenylation site. Reads with aligned 3′ ends upstream of a polyadenylation site (internal to the gene) are represented with negative distances, and downstream of a polyadenylation site (external to the gene) are represented with positive numbers. The dashed lines are at -60 and +10 from an annotated polyadenylation site, 99% of mRNA scaffolds fall within this range

**Figure 6.**
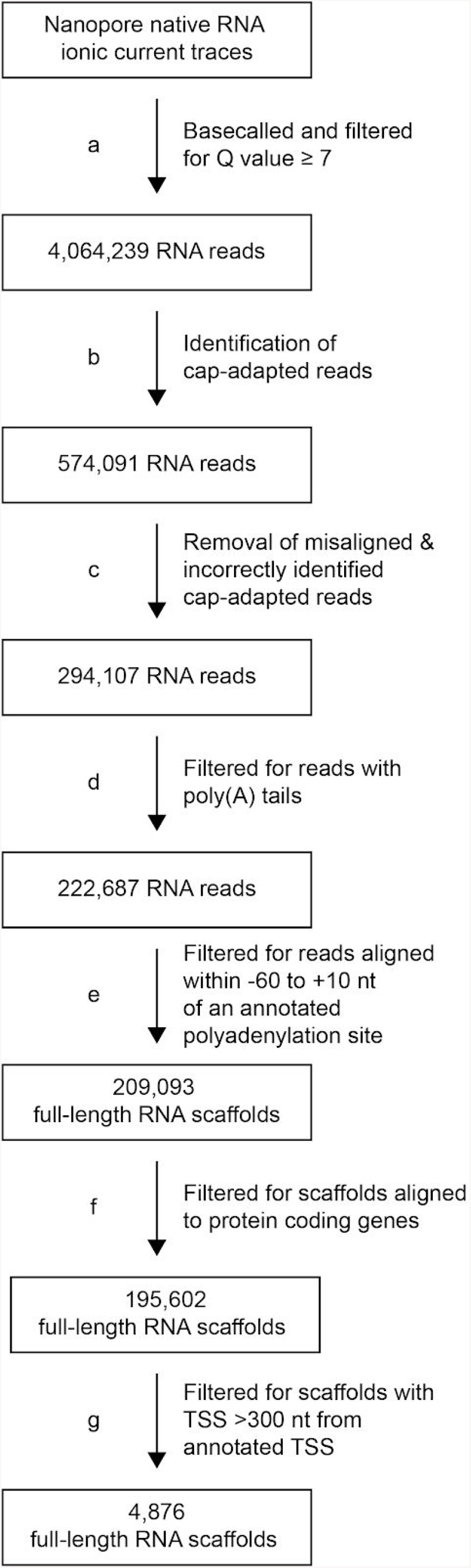
Nanopore data processing steps. Each arrow represents a computational step. Each box represents the type and quantity of data after a computational step.

We then performed a statistical measure of confidence for each full-length RNA scaffold using mapping quality scores^31^. These mapping quality scores for minimap2 range from zero (equal probability that the scaffold aligned to more than one position in the reference genome) to 60 (∼1×10^−6^ probability that the alignment was in the wrong position). Among the 209,093 RNA scaffolds, 151,182 (72.3%) had mapping quality scores of 60. There were 14,282 (6.8%) full-length RNA scaffolds with mapping quality scores of zero. Among the 4,876 full-length RNA scaffolds for unannotated isoforms, 4,568 (93.7%) had mapping quality scores of 60. There were 49 (1.0%) full-length RNA scaffolds with mapping quality scores of zero. By comparison, the untreated reads had 71.2% of the reads with a mapping quality score of 60.

### Use of high confidence scaffolds to define candidate human mRNA isoforms

We proposed that high confidence RNA scaffolds could help identify previously unannotated isoforms at sufficient precision to warrant further detailed biological experimentation. The following two examples illustrate a pipeline we used to characterize two unannotated candidate mRNA isoforms.

*Diacylglycerol O-Acyltransferase 1 (DGAT1)* encodes a multi-pass transmembrane protein that catalyzes the conversion of diacylglycerol and fatty acyl CoA to triacylglycerol. There are six annotated isoforms in GENCODE^4^ and two annotated isoforms in RefSeq^32^ (Supplementary Fig. 4). Among thirty aligned untreated nanopore reads, two reads had 5′ exons that are not documented by GENCODE^4^ nor by RefSeq^32^. In neither case was it possible to determine if the 5′ ends represented a mature mRNA transcript or a truncation product. Importantly, a single high-confidence mRNA scaffold corresponded to one of these presumptive isoforms. This confirmed connectivity between an m^7^G cap, the unannotated first exon, seventeen exons present in known isoforms, and a confirmed poly(A) tail.

*Adhesion G protein-coupled receptor E1 (ADGRE1)*. ADGRE1 is a class II adhesion GPCR that is expressed in differentiated cells in the human myeloid lineage^33^. ADGRE1 is often used as a biomarker for macrophages, however, its function is unknown^33^. The five annotated human isoforms (Fig. 7a) encode proteins with extracellular EGF-like binding domains and transmembrane domains^34^.

**Figure 7.**
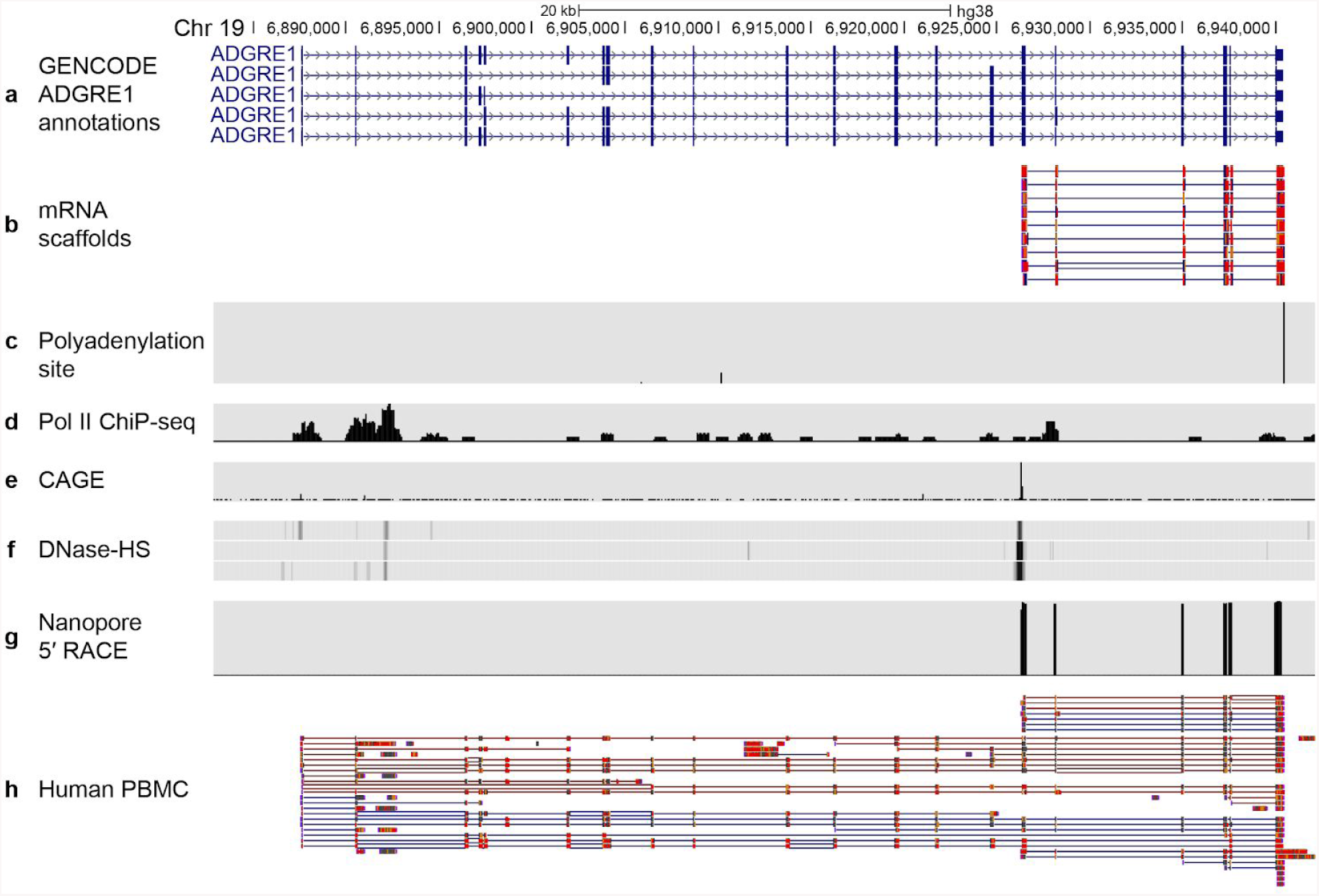
mRNA scaffolds predict an unannotated ADGRE1 isoform. (**a**) GENCODE v32 annotations for ADGRE1 mRNA isoforms. (**b**) Nine high confidence mRNA scaffolds for an unannotated ADGRE1 isoform. (**c**) Polyadenylation sites annotated by the Poly(A)Site 2.0 atlas^30^. GM12878 specific data from: (**d**) PolII ChiP-seq sites^44^; (**e**) CAGE sites for the positive strand^45^; (**f**) Three replicate DNase-HS tracks^35^. (**g**) Read coverage from full-length nanopore 5′ RACE sequencing. (**h**) Human peripheral blood mononuclear cell (PBMC) nanopore cDNA reads^36^. Red and blue lines indicated forward and reverse alignments respectively.

In our high confidence cap-adapted data set, each of nine individual RNA scaffolds aligned to a proposed ∼1,100 nt long unannotated isoform of ADGRE1 (Fig. 7b). This proposed isoform had a TSS that was internal to the annotated ADGRE1 isoforms. The scaffolds included six previously documented exons, that together encoded an in-frame ORF consistent with a protein composed of transmembrane domains 3-to-7 of the annotated ADGRE1 receptors. The extracellular amino terminus and transmembrane domains 1 and 2 were absent in the isoform predicted by the nine scaffolds.

The expected median identity is 87% for nanopore RNA sequencing reads^13^. Consequently, additional information would be needed to establish a high confidence isoform based on a single nanopore mRNA scaffold. The steps we used to substantiate the candidate ADGRE1 isoform were:

i. Confirmation that each of the nine scaffolds had a high mapping quality score (60), and that there was a poly(A) site proximal to the 3′ ends (Fig. 7c)^30^;
ii. use of orthogonal GM12878 data^35^ to support or refute the proposed isoform. We found that Pol II ChiP-seq, CAGE, and DNase-seq data all supported the proposed unannotated ADGRE1 isoform (Fig. 7d-f);
iii. 5′ RACE for the full-length proposed unannotated ADGRE1 isoform confirmed the 5′ end and revealed amplicons with identical exon composition as the RNA nanopore scaffolds (Fig. 6g);
iv. confirmation that the proposed ADGRE1 isoform was expressed in human primary tissue. It was possible that the isoform was an artifact specific to the immortalized GM12878 cell line. To rule this out, we examined long-read cDNA sequencing data from primary human peripheral blood mononuclear cells (PBMC)^36^. Seven out of ∼fifty reads that aligned to ADGRE1 were identical to the unannotated isoform identified by the nanopore scaffolds (Fig. 7h).

We also noted a consistent U-to-C miscall in the ADGRE1 stop codon of the high confidence scaffolds (Fig. 8b) and in the stop codon of all other reads from this study that aligned to ADGRE1 (Fig. 8c and d). The same pattern was observed for direct nanopore reads from a prior study that aligned to longer GENCODE annotated ADGRE1 isoforms (Fig 8e). This pattern is consistent with nanopore miscalls caused by the conversion of uridine to pseudouridine (***ψ***) at U516 in E. coli 16S rRNA^15^. Base miscalls relative to canonical training data in our ADGRE1 aligned reads (Fig. 8g) strongly suggest an unannotated pseudouridine in the stop codon, which is known to cause translation read through^37^. Importantly, this miscall was absent from *in vitro* transcribed nanopore direct RNA sequence data^11–13,15^.

**Figure 8.**
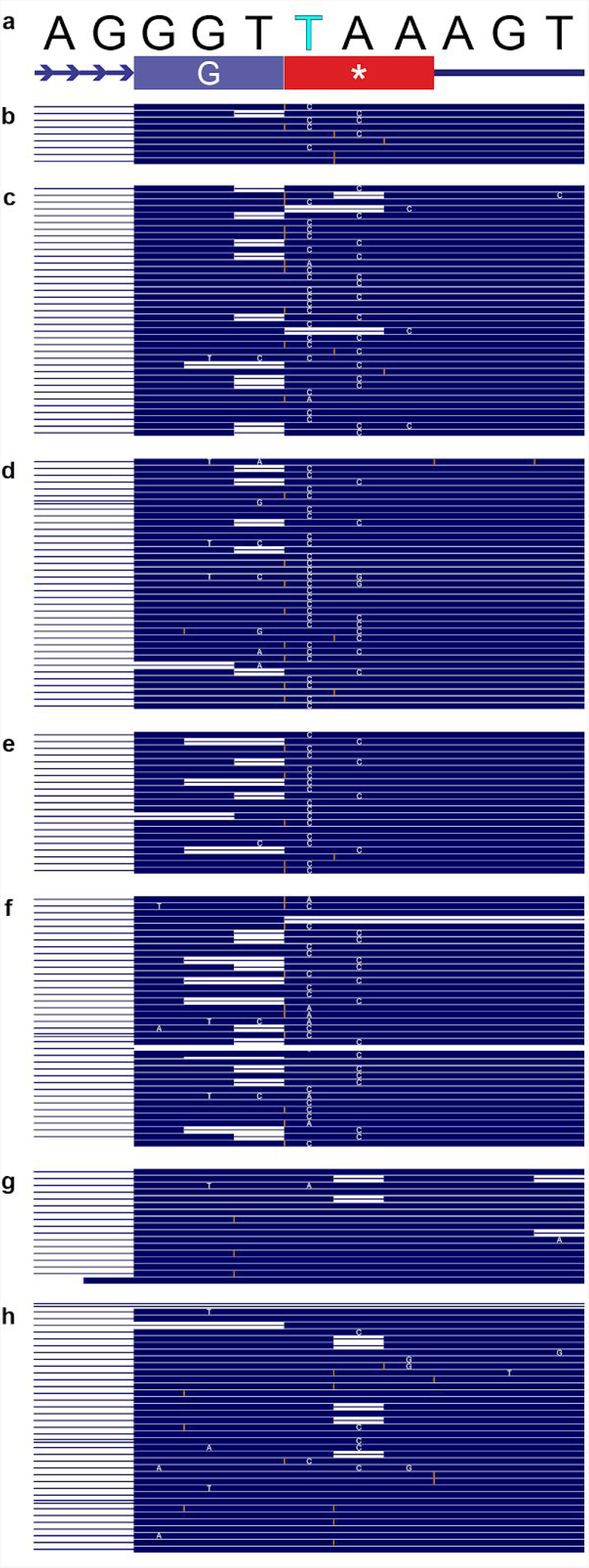
Nanopore evidence for pseudouridine in the stop codon of ADGRE1 mRNA. (**a**) HG38 chr19:6,940,022-6,940,032 which corresponds to eleven nucleotides in the last exon of ADGRE1. The G (blue background) in the second row is a glycine of the ADGRE1 gene product. The * (red background) is the canonical stop codon for ADGRE1. A pseudouridine at the first nucleotide of that stop codon can promote ribosome read through in other genes^37^. (**b**) High confidence mRNA scaffolds aligned to the unannotated isoform. (**c**) Treated-sample reads aligned to the unannotated isoform. (**d**) Untreated-sample reads aligned to the unannotated isoform. (**e**) RNA reads corresponding to an annotated ADGRE1 isoform from a previous study^13^. (**f**) RNA reads corresponding to the unannotated ADGRE1 isoform from a previous study^13^. (**g**) In vitro transcript reads composed of canonical nucleotides^13^. (**h**) cDNA amplicon reads derived from ^13^. In panels (b-to-h), dark blue is nanopore base calls that match the reference sequence. Thick white horizontal lines are nucleotide deletions in the nanopore reads. Orange vertical lines are nucleotide insertions in the nanopore reads. White letters are base calls that disagree with the reference sequence.

## DISCUSSION

In this study, we describe a strategy that uses individual nanopore reads to define high confidence human poly(A) RNA scaffolds. These scaffolds include the 5′ m^7^G cap, the 3′ polyadenylation site, and the intervening sequence at 87% identity. A majority of these scaffolds had a mapping quality score of Q60. Most of these scaffolds (95%) confirmed isoforms previously annotated in GENCODE v32^4^. There were also 4,876 full-length RNA scaffolds whose TSS were not annotated in GENCODE v32^4^, a majority of which were validated by orthogonal transcription initiation markers. Most of these TSS were internal to known mRNA isoforms.

This strategy includes a new chemo-enzymatic method to specifically adapt 5′ capped RNA strands. The RNA oligonucleotide component of the cap-adapter permitted both identification of the biological 5′ end and sequencing of approximately six additional nucleotides that were systematically missed using the conventional ONT RNA sequencing protocol. Due to the nature of the cap-adapter linkage, approximately five nucleotides are still missed at the 5′ end of each strand.

The 5′ capping procedure described in this study can attach the oligonucleotide adapter to triphosphate or diphosphate 5′ termini and to m^7^G capped 5′ termini^19^. Therefore it was conceivable that the poly(A) RNA dataset contained transcripts produced by RNA polymerases other than polII. However, when we screened for pre-processed ribosomal RNA which bears triphosphate 5′ ends, we found that only 11 transcripts out of 574,091 total cap-adapted reads aligned to ribosomal RNA genes.

In this study, we used GENCODE v32^4^ to identify annotated and unannotated isoforms in our high confidence poly(A) RNA scaffold data. This gene model set is the method of choice for high-throughput RNA analysis. RefSeq is an alternative gene model set that is often used for human genetics^32^. The unannotated ADGRE1 exemplar was absent in both gene models. However, in other cases, we found annotations in RefSeq^32^ that matched isoforms from our nanopore data that were absent in GENCODE v32^4^. Two examples are Profilin 1 (PFN1) and Voltage Dependent Anion Channel 1 (VDAC1). We recommend comparing proposed unannotated isoforms against both of these gene model sets.

We substantiated a proposed in-frame ADGRE1 isoform using four criteria that could be broadly applied. Two of these criteria were evaluated in a few hours using standard alignment visualization software (e.g. the UCSC Genome Browser) and orthogonal data for the GM12878 transcriptome. A third criterion required an additional experiment using full-length 5′ RACE amplicons that were sequenced using nanopores. This was completed in approximately three days. The fourth criterion (expression in human primary tissue) was achieved in a few days through consultation with a colleague with expertise in immune cell transcriptomics.

Unambiguous proof that this and other proposed mRNA isoforms are translated by the ribosome will require protein evidence.

We anticipate a number of improvements in the technology going forward, some of which depend on platform improvements by ONT. For example, the percent identity of ONT direct RNA base calls has remained at ∼87% since the technology was introduced in 2016. By comparison, the percent identity for DNA base calls has increased from ∼66% in 2014^38^ to ∼96.4% in 2020 (unpublished UCSC data). Other improvements could be implemented by the research community. For instance, preliminary data suggest that the capping efficiency can approach ∼84% on synthetic RNA (data not shown). This suggests that the percent of cap-adapted reads identified by nanopore sequencing could substantially improve from the 14.3% observed in this study. Furthermore, the 5′ end adapter used in this study included a PEG spacer that causes the ONT motor enzyme to slip and thus miss ∼five nucleotides at the 5′ end of each strand. An adapter structure that allowed for sequencing of those five nucleotides would be useful, especially because the N1 and N2 positions are often modified^16^.

In summary, NRCeq enabled identification of individual, high-confidence, RNA scaffolds representing annotated and unannotated full-length human RNA isoforms.

## ONLINE METHODS

### Synthesis of 3′-DBCO RNA adapter

The 45-nucleotide 3′-DBCO RNA oligomer (CUCUUCCGAUCUACACUCUUUCCCUACACGACGCUCUUCCGAUCU) was synthesized by coupling the 3′-NH_2_ RNA oligomer with a DBCO-sulfo-NHS ester (Glen Research, #50-1941). The 3′-NH_2_ RNA synthesis was performed on an ABI 394 DNA synthesizer (Applied Biosystems) starting with 3′-PT-amino-modifier C3 CPG (Glen Research, #20-2954) and UltraFast RNA TBDMS RNA amidites (Glen Research: Bz-A-CE #10-3003, Ac-C #10-3015, Ac-G-CE #10-3025, and U-CE #10-3030). The oligonucleotide was deprotected according to the manufacturer’s protocol using ammonium hydroxide/methylamine and purified using a Glen-Pak RNA purification cartridge (Glen Research, #60-6100) followed by PAGE. The purified 3′-NH_2_ RNA was dissolved in 5 mL of 0.1 M sodium borate (pH 8.3). Then 2.5 mL of a 20 mM solution of DBCO-sulfo-NHS ester in DMSO was added and stirred for 1.5 h at room temperature. The reaction was then dissolved in 0.1 M TEAB (up to 35 mL) and purified by C8 RP-HPLC (Higgins Analytical) using 0.1 M TEAB and acetonitrile as mobile phase. The 3′-DBCO RNA oligonucleotide was concentrated and re-purified by PAGE and desalted using a Clarity-RP desalting cartridge (Phenomenex, #8B-S041-HBJ).

### 5′ RACE library

Total RNA from GM12878 (12 µg) was treated with QuickCIP (NEB, #M0525) at 0.5 U/µL in the provided buffer at 37 °C for 20 min and then purified using RNA Clean & Concentrator (Zymo Research, #R1013) with the standard protocol. The RNA was divided into two aliquots (1 and 2). Aliquot 1 was treated with RppH (NEB, #M0356) in 1X Thermopol Buffer (NEB, #B90004) at 0.5 U/µL at 37 °C for 1 h, whereas aliquot 2 was incubated under the same conditions in the absence of RppH to serve as a non-decapped control. The two samples were purified as above and eluted in 20 µL of RNase-free water. Each sample (10 µL) was ligated to 5 pmol of a single-stranded RNA adapter (see below) using 1.5 µL T4 RNA Ligase I (NEB, #M0204) in 1X T4 RNA Ligase Reaction Buffer in 20 µL total volume for 1 h at 25 °C. Sample 1 (RppH treated) was ligated to the adapter SRGAUUA and sample 2 (mock treated) to adapter SRAUCAG, wherein SR denotes the sequence: GUUCAGAGUUCUACACUCCGACGAUC. The 3′ terminal five nucleotides of each adapter served as an identification index for the provenance of the sequence products (sample 1, decapped vs. sample 2, not decapped). After ligation the two samples were pooled and mixed with 80 µL AMPure XP magnetic beads (Beckman Coulter) and processed for magnetic purification of the RNA, which was eluted in 20 µL water. The RNA was used for first strand cDNA synthesis with random priming using the Protoscript II First strand synthesis kit (NEB, #E6560) in a total volume of 40 µL for 1 h at 42 °C. The cDNA was diluted to 150 µL and aliquoted into 96 wells of a PCR plate (1.5 µL/well) each of which contained 20 pmol of a reverse primer specific for each target sequence (see Table X for target genes and primer sequences). The PCR reactions were performed using the forward primer AATGATACGGCGACCACCGAGATCTACACGTTCAGAGTTCTACAGTCCGA, and LongAmpTaq (NEB, #M0287) with the following program: 94 °C 1 min, followed by 5 cycles of 94 °C 10 s, 60 °C 15 s, 65 °C 15 s, followed by 32 cycles of 94 °C 10 s, 55 °C 15 s, 65 °C 15 s, followed by 65 °C 5 min. An aliquot from each reaction (2 µL) reaction was evaluated for quality and to estimate product concentration using the Agilent 2200 Tapestation system. A fraction of each reaction (varying from 1-10 µL depending on concentration) was used for pooling to obtain a mixture of all 96 PCR products. The DNA mixture was purified using 200 µL AMPure XP beads per 125 µl DNA, eluted in 40 µL water. Illumina adapters were added by amplifying the sample for 4 cycles using the SR primer and index primer from the kit (NEB, #E7330). The resulting product was purified with AMPure XP beads at 1:1 ratio, and the eluted material was sequenced in an Illumina Miseq sequencer using paired end 2×150 nt. The FASTQ files were processed to identify the 5′ end terminal sequence of each targeted transcript. RppH treated and untreated reads were distinguished using the cutadapt (version 2.10) with the following parameters : cutadapt -O 5 --action lowercase --trimmed-only --pair-filter first and -g ^GATTA (for reads from the RppH treated sample) and -g ^ATCAG (for reads from the untreated sample). The resulting paired end reads were trimmed using Cutadapt wrapped with trim galore (version 0.3.3)^39^. The trimmed paired-end reads were aligned to the human genome using STAR_2.4.0b^40^ using default parameters and the mapped read 2 was discarded. The ratio of the number of R1 reads from the RppH treated sample over the sum of reads from both samples mapping within 100 bp window of a predicted TSS position was calculated. For a TSS to be considered validated, a ratio higher than 0.5 was required with at least one RACE read from the RppH treated sample.

### GM12878 cell tissue culture

GM12878 cells were cultured the same as in ^13^. Briefly, GM12878 cells (passage 11) were cultured in RPMI medium (Gibco, #21870076) supplemented with 15% non heat-inactivated FBS (Gibco, #12483020) and 2 mM L-Glutamax (Gibco, #35050061). Cells were expanded to 9 × T75 flasks (45 mL of medium in each) and centrifuged for 10 min at 100*g* (4 °C), washed in 1/10th volume of PBS (pH 7.4), and combined for homogeneity. The cells were then evenly split between 8 × 15 mL tubes and pelleted at 100*g* for 10 min at 4 °C. The cell pellets were then snap frozen in liquid nitrogen and immediately stored at −80 °C.

### Isolation of GM12878 total RNA

GM12878 RNA was isolated the same as in ^13^. Briefly, 4 mL of TRI-Reagent (Invitrogen, #AM9738) was added to a frozen pellet of 5 × 10^7^ GM12878 cells and vortexed immediately. This sample was incubated at room temperature for 5 min. CHCl_3_ (chloroform, 200 µL) was added per mL of sample, vortexed, incubated at room temperature for 5 min, vortexed again, and centrifuged for 10 min at 12,000*g* (4 °C). The aqueous phase was pooled in a LoBind Eppendorf tube and combined with an equal volume of isopropanol. The tube was mixed, incubated at room temperature for 15 min, and centrifuged for 15 min at 12,000*g* (4 °C). The supernatant was removed, the RNA pellet was washed with 750 μL 80% ethanol and then centrifuged for 5 min at 12,000*g* (4 °C). The supernatant was removed. The pellet was air-dried for 10 min, resuspended in nuclease-free water (100 μL final volume), quantified, and either stored at –80 °C or processed further by Poly(A) purification.

### GM12878 poly(A) RNA purification

Poly(A) RNA was purified from GM12878 total RNA with NEXTflex poly(A) beads (Bioo Scientific, NOVA-512980) using 50 µL of beads per 100 µg of total RNA. GM12878 poly(A) RNA was aliquoted and stored at –80°C.

### Isolation of *S. cerevisiae* S288C total RNA

Total RNA was purified from Saccharomyces cerevisiae S288C. The *S. cerevisiae* was grown in 1 L YPD media (1% yeast extract, 2% peptone, 2% dextrose) at 30 °C. The cells were pelleted and resuspended in cold 10 mM EDTA. The cells were again pelleted and resuspended in 5 mL of 50 mM sodium acetate (pH 5.5), 10 mM EDTA, 1% SDS. 5 mL of Acid-Phenol:Chloroform:IAA (Invitrogen, #AM9720) was added, and the mixture was vortexed. The mixture was incubated in a 65 °C water bath with brief vortexing every 5 min for a total incubation time of 30 min. The mix was placed on ice for 10 min, and the phases separated by centrifugation. The upper phase was collected, and an equal volume of chloroform was added. The mixture vortexed again, and the phases separated by centrifugation. The upper phase collected and 0.1 volume of 3 M sodium acetate pH 5.3 was added. An equal volume of isopropanol was added, mixed, and the RNA was precipitated at –20 °C. The resulting RNA precipitate was dissolved in 5 mL of TE buffer. The RNA was reprecipitated by adding 0.25 volume of 1 M sodium acetate pH 5.5 and 2.5 volumes of ethanol and incubated for 60 min at –20 °C. The total RNA pellet was redissolved in TE buffer.

### *S. cerevisiae* S288C poly(A) RNA purification

Poly(A) RNA was isolated from 2 mg of total *S. cerevisiae* RNA using the PolyA Spin mRNA Isolation Kit (NEB, #S1560). After a single round of isolation, the RNA was precipitated by adding glycogen and 2.5 volumes of ethanol. The poly(A) RNA pellet was dried and resuspended in 1 mM Tris-HCl pH 7.5, 0.1 mM EDTA.

### Decapping and recapping of poly(A) RNA

Poly(A) RNA was decapped and recapped according to methods previously described ^21^. In brief, decapping of 1.5-6 µg poly(A) RNA was performed with 1.5 µL yDcps (NEB, #M0463) in 1X yDcpS reaction buffer (10 mM Bis-Tris-HCl pH 6.5, 1 mM EDTA) in 50 µL total volume for 1 h at 37 °C. The decapped RNA was purified using RNA Clean and Concentrator (Zymo Research, #R1013) with the standard protocol (>17 nt recovery) and eluted in 30 µL of RNase-free water. Recapping the 5′ end of the decapped poly(A) RNA was performed with 6 µL *Vaccinia* Capping Enzyme (VCE) (NEB, #M2080) in 1X VCE reaction buffer (50 mM Tris HCl, 5 mM KCl, 1 mM MgCl2, 1 mM DTT, pH 8), 6 µL *E. coli* Inorganic Pyrophosphatase (NEB, #M0361), 0.5 mM 3′-azido-ddGTP (Trilink, #N-4008), 0.2 mM *S*-adenosylmethionine (SAM) (NEB, #B9003) in 60 µL total volume for 30 min at 37 °C. The recapped RNA was purified with RNA Clean and Concentrator as above.

### Adaptation of recapped poly(A) RNA

Azido-ddGTP recapped RNA (1-2 µg) was concentrated briefly on a SpeedVac vacuum concentrator (Savant) to reduce the volume to approximately 5-10 µL. Copper-free Click Chemistry reactions were performed in a total volume of 50 μL, containing 25% v/v PEG 8000 (NEB, #B1004) and 20% v/v acetonitrile (Sigma-Aldrich, #271004) in 0.1 M sodium acetate buffer, pH 4 (10X, Alfa Aesar, #J60104) and 10 mM EDTA (50x, Invitrogen, #15575-038). Azido-ddGTP recapped RNA and the 3′-DBCO RNA adapter (200 nmol, final concentration of 4 µM) were added and shaken for 2 h at room temperature. Then, acetonitrile was removed by brief concentration on a SpeedVac, and the adapted RNA recovered using RNA Clean & Concentrator (Zymo Research, #R1013) following the protocol to separate large RNA (desired) from small RNA (excess adapter).

### Validation of an unannotated ADGRE1 isoform

cDNA for 5′ RACE sequencing was made with the 5′ RACE Protocol using the Template Switching RT Enzyme Mix (NEB, #M0466).A ADGRE1 cDNA was reverse transcribed from total GM12878 RNA using a template-switching oligo (TSO) (GCTAAT<u>CATTGCAAGCAGTGGTATCAAC</u>GCAGAGTACATrGrGrG) a poly(dT) reverse transcription primer. ADGRE1 cDNA was PCR amplified using a forward primer (underlined sequence of the TSO) and a gene-specific reverse primer with Q5 Hot Start High-Fidelity 2X Master Mix (NEB, #M0494S). cDNA was prepared for sequencing using the barcoded NBD 104 expansion of the SQK-LSK109 protocol following the manufacturer’s recommendations and sequenced using a Flongle flow cell. Ionic current traces were basecalled with MinKnow real-time basecalling using the high-accuracy model.

### MinION RNA sequencing

Poly(A) RNA samples were split and either processed for cap-adaptation (treated) or used as a matched negative control (untreated). Both treated and untreated poly(A) RNA (500-775 ng) were prepared for nanopore direct RNA sequencing generally following the ONT SQK-RNA002 kit protocol, including the optional reverse transcription step recommended by ONT. Instead of using Superscript III, as in the ONT protocol, Superscript IV (Thermo Fisher, #18091050) was used for reverse transcription. RNA sequencing on the MinION was performed using ONT R9.4 flow cells and the standard MinKNOW protocol (48 h sequencing script) as recommended by ONT, with one exception. We collected bulk phase continuous data files for 2 h of sequencing and then restarted the sequencing runs after the two hours of initial sequencing.

### Basecalling, filtering and alignments

We used the ONT Guppy workflow (version 3.0.3+7e7b7d0 configuration file rna_r9.4.1_70bps_hac.cfg) for basecalling direct RNA. NanoFilt (version 2.5.0)^41^ was used to classify reads as pass if the pre-read average Phred-score threshold was greater than or equal to 7 and fail if less than 7. A custom python script was used to convert the U’s in the guppy basecalled sequence to T’s. Porechop (version 0.2.4) was used to identify the 5′ adapter sequence (https://github.com/rrwick/Porechop). We used barcode_diff 1 and barcode_threshold 70 or 74 for S288C or GM12878 reads, respectively. The adapters were untrimmed while optimizing parameters and trimmed for all other analysis. The barcode search sequence, TCCCTACACGACGCTCTTCCGA, was added to the end of the adapter list in the adapters.py. Reads were then aligned to the appropriate reference, sacCer3 or GRCh38, using minimap2^42^ (version 2.16-r922) with recommended conditions.

### TSS filtering pipeline

TSS filtering pipeline is available on github (https://github.com/mitenjain/dRNA_capping_analysis). Reads were mapped to the human genome (GRCh38.p3.genome.fa) using Minimap2 (version 2-2.9) with the following parameters : --secondary=no -ax splice -k14 -uf. Remaining secondary alignments were removed (using samflag -F 2048). Reads containing 15 or more soft or hard clipped bases at the 5′end of the read were removed to avoid positioning TSS at the wrong locations (properly mapped reads). Finally, for adapted reads only, the untrimmed reads containing less than 15 soft or hard clipped bases were flagged and the equivalent trimmed reads removed. This filter removes the reads that were called adapted by poreshop but the adapter is matching the genomic sequence (true positive reads). Unless otherwise stated, subsequent analyses for the untreated sample or unadapted fraction of the treated sample were done using the properly mapped reads. For the adapted fraction of the treated sample, subsequent analysis was done on the properly mapped/true positive reads.

### Identification of novel 5′ends

In order to identify TSS that have not previously been annotated, only the 5′end reads mapping to 300 or more bases away from a GENCODE v32^4^ annotated TSSs were retained. Next, we used reads that aligned to GENCODE genes. Default parameters were used to filter the GM12787 data. GENCODE v32^4^ was used as our known isoform and start site annotation.

### 5′ RACE candidate gene selection

RACE candidates were selected from the unannotated genes identified by the TSS filtering pipeline. Genes with a cap-adapted read which had a TSS at unannotated exons or internal exons were considered as candidates. Transcripts with a longer or shorter annotated first exon were excluded from this analysis. 88 candidate genes were selected for 5′ RACE validation.

### Mapping 5′ end to chromatin marks, ChIP-seq and CAGE data

Encode data were downloaded using the following accession numbers : ENCFF340BYJ (POLR2 ChIP-seq), ENCFF289XSX (SPI1 ChIP-seq), ENCFF093VXI (DNase-seq) and ENCFF580WIH (CAGE). For plotting purposes, all the datasets were downsampled to the total number of read defining novel TSS in the treated sample (9116 reads). To avoid repeats in the dataset, all the reads within a 50 bp window were merged. The most 5′end of the mapped reads were used to define the reference locations. Deeptools 3.3.0^43^ was used to perform the heatmap plotting with the following parameters : computeMatrix reference-point -a 1000 -b 1000 and plotHeatmap --missingDataColor=#440154FF --colorList ‘#440154FF,#238A8DFF,#FDE725FF’ ‘#440154FF,#238A8DFF,#FDE725FF’ ‘#440154FF,#238A8DFF,#FDE725FF

### Annotation of the nanopore reads

The genomic position of the 5′ end of mapped nanopore reads was compared to GENCODE annotation v32^4^ using bedtools (v2.27.1) with the following parameters : bedtools closest -t first -D a -iu -s. For each matching annotation, the gene_type information was used to quantify the overlap with a specific gene type (for example “protein-coding”). The quantification was normalized to 1 and plotted in a bar plot with colors representing the gene_type.

## Data and code availability

Data and analysis scripts can be found at the following github, https://github.com/mitenjain/dRNA_capping_analysis.

## Acknowledgments

Guillermo Chacaltana generated the nanopore full-length ADGRE1 cDNA data. Chris Vollmers provided the long-read PBMC cDNA data. Jonathan Mudge and Irwin Jungreis gave advice on GENCODE annotations. Kristof Tigyi cultured the GM12878 cells. Niki Thomas and Robin Abu-Shumays edited drafts of the manuscript. Larry McReynolds and Bill Jack gave helpful comments. We acknowledge the ENCODE Consortium and the following encode data producers for their assistance: J. Michael Cherry, Stanford; Gregory Crawford, Duke; and John Stamatoyannopoulos, University of Washington. We downloaded call sets from the ENCODE portal^35^ (https://www.encodeproject.org) with the following identifiers: ENCFF743ULW, ENCFF093VXI, ENCFF066VBS, and ENCFF969DFL, ENCSR000EMT, ENCSR000EJD, ENCSR000EMT, ENCSR000CKA, ENCSR000CJZ, ENCSR000CJY. This work was supported by New England Biolabs Inc. and the following grants: NIH HG010053 (M.A.), Oxford Nanopore Research Grant SC20130149 (M.A.), NHGRI U54HG004555 (M.D.), and gifts from the W. H. Akeson Research Fund.

