## SupplementalData for "Identification of high confidence human poly(A) RNA isoform scaffolds using nanopore sequencing"

### SUPPLEMENTARY INFORMATION

#### SUPPLEMENTARY TABLES

**Supplementary Table 1** Native RNA nanopore sequencing statistics

| Organism | sample | "Click" type | pass reads | N50 | adapted |
|---|---|---|---|---|---|
| S. cerevisiae | Untreated 1 | None | 1,391,578 | 531 | 0.0000 |
| S. cerevisiae | Untreated 2 | None | 1,755,395 | 1,092 | 0.0000 |
| S. cerevisiae | Untreated 3 | None | 3,222,192 | 1,034 | 0.0000 |
| S. cerevisiae | Untreated 4 | None | 861,210 | 947 | 0.0000 |
| **S. cerevisiae** | **Untreated Pooled** | **None** | **7,230,375** | **957** | **0.0000** |
| S. cerevisiae | Treated 1 | $Cu^{2+}$ | 2,576,122 | 798 | 0.2382 |
| S. cerevisiae | Treated 2 | $Cu^{2+}$ | 3,933,777 | 676 | 0.0889 |
| S. cerevisiae | Treated 3 | $Cu^{2+}$ | 1,848,281 | 722 | 0.1451 |
| S. cerevisiae | Treated 4 | $Cu^{2+}$ | 1,382,164 | 575 | 0.0696 |
| S. cerevisiae | Treated 5 | $Cu^{2+}$ | 600,225 | 435 | 0.0618 |
| **S. cerevisiae** | **Treated Pooled** | **$Cu^{2+}$** | **10,340,569** | **692** | **0.1340** |
| S. cerevisiae | Treated 1 | Cu-Free | 1,128,595 | 737 | 0.3347 |
| S. cerevisiae | Treated 2 | Cu-Free | 3,799,042 | 755 | 0.4138 |
| S. cerevisiae | Treated 3 | Cu-Free | 1,275,112 | 715 | 0.3407 |
| **S. cerevisiae** | **Treated Pooled** | **Cu-Free** | **6,202,749** | **744** | **0.3844** |
| GM12878 | Untreated 1 | None | 1,332,182 | 1,572 | 0.0000 |
| GM12879 | Untreated 2 | None | 2,497,808 | 1,691 | 0.0000 |
| **GM12880** | **Untreated Pooled** | **None** | **3,829,990** | **1,615** | **0.0000** |
| GM12881 | Treated 1 | Cu-Free | 640,221 | 1,334 | 0.1106 |
| GM12882 | Treated 2 | Cu-Free | 919,701 | 1,131 | 0.1021 |

| | | | | | |
|---|---|---|---|---|---|
| GM12883 | Treated 3 | Cu-Free | 797,451 | 1,212 | 0.1596 |
| GM12884 | Treated 4 | Cu-Free | 1,706,866 | 1,189 | 0.1653 |
| **GM12885** | **Treated Pooled** | **Cu-Free** | **4,064,239** | **1,207** | **0.1413** |

**Supplementary Table 2** Effect of copper-catalyzed and copper-free click reactions on RNA integrity and nanopore read quality. The RIN was measured from S. cerevisiae total RNA after enzyme treatment and purification for each step of the cap-adaptation process using an Agilent RNA 6000 Nano Kit (mean ± SD for n = 2 experiments). Percent cap-adapted is the percent of poly(A) RNA nanopore reads identified by Porechop as cap-adapted. The read N50 is where half of the total bases sequenced are in reads of that length or longer.

| | **No Treatment** | **yDCPS** | **VCE** | **Copper-catalyzed Click Adaptation** | **Copper-Free Click Adaptation** |
|---|---|---|---|---|---|
| **RIN** | 9.5 ± 0.1 | 9.4 ± 0.4 | 8.1 ± 0.2 | 6.7 ± 0.2 | 8.1 ± 0.7 |
| **Percent cap-adapted** | - | - | - | 13.4% | 38.4% |
| **N50** | 957 nt | - | - | 692 nt | 744 nt |

**Supplementary Table 3** Counts of RNA types for untreated, unadapted, and cap-adapted reads.

| RNA type | untreated | unadapted | cap-adpated |
|---|---|---|---|
| protein_coding | 988063 | 1954644 | 278718 |
| Mt_tRNA | 59686 | 110205 | 6 |
| processed_pseudogene | 26917 | 59970 | 7014 |
| Mt_rRNA | 26325 | 39885 | 154 |
| lncRNA | 22023 | 38564 | 4642 |
| unknown | 20129 | 35120 | 2823 |
| transcribed_processed_pseudogene | 4011 | 9612 | 1447 |
| transcribed_unprocessed_pseudogene | 3035 | 6087 | 631 |
| IG_V_gene | 1540 | 1818 | 1087 |
| unprocessed_pseudogene | 670 | 1463 | 189 |
| IG_C_gene | 554 | 6050 | 26 |
| polymorphic_pseudogene | 317 | 905 | 426 |
| snRNA | 271 | 93 | 0 |
| miRNA | 252 | 549 | 37 |
| IG_V_pseudogene | 227 | 1027 | 112 |
| TR_J_gene | 175 | 117 | 81 |
| transcribed_unitary_pseudogene | 172 | 395 | 31 |
| rRNA | 157 | 91 | 2 |
| snoRNA | 143 | 104 | 0 |
| TEC | 128 | 249 | 35 |
| misc_RNA | 91 | 69 | 16 |
| TR_C_gene | 64 | 252 | 2 |

| | | | |
|---|---|---|---|
| IG_J_gene | 52 | 433 | 12 |
| unitary_pseudogene | 14 | 25 | 1 |
| snoRNA | 5 | 3 | 8 |
| translated_processed_pseudogene | 5 | 7 | 1 |
| TR_V_gene | 4 | 6 | 1 |
| scRNA | 3 | 2 | 0 |
| TR_V_pseudogene | 3 | 2 | 1 |
| translated_unprocessed_pseudogene | 3 | 8 | 6 |
| IG_J_pseudogene | 2 | 15 | 1 |
| TR_D_gene | 2 | 0 | 0 |
| TR_J_pseudogene | 2 | 0 | 0 |
| snRNA | 1 | 2 | 16 |

**Supplementary Table 4** Nanopore Recappable-seq TSS validation by 5′ RACE. 90 genes comprising 95 TSS (including two control genes) are listed below. 5′ RACE signal validated 64 TSS for 61 genes. Weak support indicates a low number of reads.

| Gene | Chromosome | start | end | strand | RPPH reads | no RPPH reads | RPPH % | CONFIRMED | Comments |
|---|---|---|---|---|---|---|---|---|---|
| AMBRA1 | chr11 | 46483098 | 46483198 | - | 3442 | 0 | 100 | YES | |
| BCL2A1 | chr15 | 80017734 | 80017834 | - | 7392 | 0 | 100 | YES | |
| BLK | chr8 | 11538185 | 11538285 | + | 1115 | 0 | 100 | YES | |
| CCDC12 | chr3 | 46927226 | 46927326 | - | 3276 | 0 | 100 | YES | |
| CCR10 | chr17 | 42682390 | 42682490 | - | 147 | 0 | 100 | YES | |
| CORO1B | chr11 | 67438647 | 67438747 | - | 1 | 0 | 100 | YES | weak support |
| CRELD2 | chr22 | 49925093 | 49925193 | + | 2453 | 0 | 100 | YES | |
| FBXL15 | chr10 | 102421091 | 102421191 | + | 1625 | 0 | 100 | YES | |
| FHL2 | chr2 | 105373930 | 105374030 | - | 2001 | 0 | 100 | YES | |
| FLOT2 | chr17 | 28896992 | 28897092 | - | 1015 | 0 | 100 | YES | |
| GEMIN7 | chr19 | 45075638 | 45075738 | + | 31 | 0 | 100 | YES | |
| ICA1 | chr7 | 8134516 | 8134616 | - | 2234 | 0 | 100 | YES | 1 of 3 TSS |
| KDM4B | chr19 | 5131088 | 5131188 | + | 1 | 0 | 100 | YES | weak support |
| Laptm5 | chr1 | 30758842 | 30758942 | - | 1132 | 0 | 100 | YES | 1 of 2 TSS |
| Laptm5 | chr1 | 30746932 | 30747032 | - | 170 | 0 | 100 | YES | 1 of 2 TSS |
| MBD2 | chr18 | 54219731 | 54219831 | - | 7993 | 0 | 100 | YES | |
| MMAA | chr4 | 145625973 | 145626073 | + | 1334 | 0 | 100 | YES | |
| NFKBIE | chr6 | 44263240 | 44263340 | - | 4114 | 0 | 100 | YES | 1 of 2 TSS |
| PAGR1 | chr16 | 29816457 | 29816557 | + | 648 | 0 | 100 | YES | |
| PGLYRP4 | chr1 | 153343782 | 153343882 | - | 1888 | 0 | 100 | YES | |
| PHPT1 | chr9 | 136847953 | 136848053 | + | 294 | 0 | 100 | YES | |

| | | | | | | | | | |
|---|---|---|---|---|---|---|---|---|---|
| PLLP | chr16 | 57260595 | 57260695 | - | 33538 | 0 | 100 | YES | |
| PYROXD2 | chr10 | 98391150 | 98391250 | - | 5340 | 0 | 100 | YES | |
| SPIB | chr19 | 50419764 | 50419864 | + | 2396 | 0 | 100 | YES | |
| STAC3 | chr12 | 57249092 | 57249192 | - | 163 | 0 | 100 | YES | |
| STAG3 | chr7 | 100210960 | 100211060 | + | 880 | 0 | 100 | YES | |
| UNC13C | chr15 | 54583777 | 54583877 | + | 3811 | 0 | 100 | YES | 1 of 2 TSS |
| WDR91 | chr7 | 135188773 | 135188873 | - | 142 | 0 | 100 | YES | |
| GMNN | chr6 | 24779789 | 24779889 | + | 1780 | 1 | 100 | YES | |
| KIFAP3 | chr1 | 170071492 | 170071592 | - | 17114 | 15 | 100 | YES | |
| BAG2 | chr6 | 57172604 | 57172704 | + | 27958 | 48 | 100 | YES | |
| GPR15 | chr3 | 98536538 | 98536638 | - | 2731 | 6 | 100 | YES | |
| TPRG1 | chr3 | 189308204 | 189308304 | `+ | 31247 | 100 | 100 | YES | |
| MAP7D2 | chrX | 20056827 | 20056927 | - | 18778 | 103 | 99 | YES | |
| NDUFAF4 | chr6 | 96897383 | 96897483 | - | 6281 | 53 | 99 | YES | |
| SERINC2 | chr1 | 31423625 | 31423725 | + | 12442 | 105 | 99 | YES | |
| SMG9 | chr19 | 43748774 | 43748874 | - | 16446 | 212 | 99 | YES | |
| SUPV3L1 | chr10 | 69202538 | 69202638 | + | 1478 | 22 | 99 | YES | |
| DHRS7 | chr14 | 60210465 | 60210565 | - | 6814 | 189 | 97 | YES | |
| MFSD14A | chr1 | 100077016 | 100077116 | + | 1161 | 39 | 97 | YES | |
| NSMCE1 | chr16 | 27243237 | 27243337 | - | 1676 | 59 | 97 | YES | |
| JUP | chr17 | 41771828 | 41771928 | - | 640 | 27 | 96 | YES | |
| DENND6B | chr22 | 50314503 | 50314603 | - | 1203 | 66 | 95 | YES | |
| ICA1 | chr7 | 8128778 | 8128878 | - | 7951 | 454 | 95 | YES | 1 of 3 TSS |
| TMSB10 | chr2 | 84905617 | 84905717 | + | 315135 | 19612 | 94 | YES | control |
| 17ORF49 | chr17 | 7015451 | 7015551 | + | 31898 | 2012 | 94 | YES | |

| | | | | | | | | | |
|---|---|---|---|---|---|---|---|---|---|
| TIMD4 | chr5 | 156922249 | 156922349 | - | 47152 | 3290 | 93 | YES | |
| ACTB | chr7 | 5530538 | 5530638 | - | 15718 | 1321 | 92 | YES | control |
| MARC2 | chr1 | 220770439 | 220770539 | + | 1641 | 141 | 92 | YES | |
| ADGRE1 | chr19 | 6926293 | 6926393 | + | 22615 | 1981 | 92 | YES | |
| ELMO1 | chr7 | 36951258 | 36951358 | - | 112177 | 11189 | 91 | YES | |
| UPB1 | chr22 | 24503394 | 24503494 | + | 97503 | 10501 | 90 | YES | |
| PTPN6 | chr12 | 6954041 | 6954141 | + | 3993 | 698 | 85 | YES | |
| DOCK2 | chr5 | 169746401 | 169746501 | + | 20428 | 3811 | 84 | YES | |
| TEKT4 | chr2 | 94873323 | 94873423 | + | 2148 | 446 | 83 | YES | |
| TYMP | chr22 | 50526939 | 50527039 | - | 97 | 21 | 82 | YES | |
| CTSH | chr15 | 78939164 | 78939264 | - | 1373 | 428 | 76 | YES | |
| MSC | chr8 | 71843775 | 71843875 | - | 2595 | 826 | 76 | YES | |
| CD27 | chr12 | 6450337 | 6450437 | + | 568 | 183 | 76 | YES | |
| DR1 | chr1 | 93346386 | 93346486 | + | 4733 | 1858 | 72 | YES | |
| PLEK | chr2 | 68388968 | 68389068 | + | 77406 | 31720 | 71 | YES | |
| NFKBIE | chr6 | 44260521 | 44260621 | - | 507 | 225 | 69 | YES | 1 of 2 TSS |
| TNFRSF4 | chr1 | 1213035 | 1213135 | - | 279 | 147 | 65 | YES | |
| ANXA6 | chr5 | 151124311 | 151124411 | - | 22068 | 13327 | 62 | YES | |
| ANXA11 | chr10 | 80201540 | 80201640 | - | 1496 | 1277 | 54 | YES | |
| MAP2k2 | chr19 | 4115016 | 4115116 | - | 16 | 15 | 52 | YES | weak support |
| CD19 | chr16 | 28932342 | 28932442 | + | 9 | 43 | 17 | NO | |
| MRPS17 | chr7 | 55953127 | 55953227 | + | 96 | 1579 | 6 | NO | |
| AICDA | chr12 | 8611097 | 8611197 | - | 0 | 0 | 0 | NO | |
| ASAP1 | chr8 | 130358572 | 130358672 | - | 0 | 0 | 0 | NO | |
| ASCL1 | chr12 | 102958440 | 102958540 | + | 0 | 0 | 0 | NO | |

| | | | | | | | | | |
|---|---|---|---|---|---|---|---|---|---|
| BATF | chr14 | 75515759 | 75515859 | + | 0 | 0 | 0 | NO | |
| BBC3 | chr19 | 47228468 | 47228568 | - | 0 | 0 | 0 | NO | |
| BCHE | chr3 | 165830479 | 165830579 | - | 0 | 150 | 0 | NO | |
| CD70 | chr19 | 6590688 | 6590788 | - | 0 | 0 | 0 | NO | |
| CETP | chr16 | 56971286 | 56971386 | + | 0 | 0 | 0 | NO | |
| CNFN | chr19 | 42387393 | 42387493 | - | 0 | 0 | 0 | NO | |
| COG4 | chr16 | 70483536 | 70483636 | - | 0 | 0 | 0 | NO | |
| ECHS1 | chr10 | 133373663 | 133373763 | - | 0 | 0 | 0 | NO | |
| ENTPD2 | chr9 | 137050661 | 137050761 | - | 0 | 0 | 0 | NO | |
| FAM78A | chr9 | 131278122 | 131278222 | - | 0 | 0 | 0 | NO | |
| ICA1 | chr7 | 8176394 | 8176494 | - | 0 | 598 | 0 | NO | 1 of 3 TSS |
| IRF2BP2 | chr1 | 234610120 | 234610220 | - | 0 | 0 | 0 | NO | |
| MICAL1 | chr6 | 109454191 | 109454291 | - | 0 | 0 | 0 | NO | |
| MRPS26 | chr20 | 3046308 | 3046408 | + | 0 | 0 | 0 | NO | |
| MYBL2 | chr20 | 43710949 | 43711049 | + | 0 | 0 | 0 | NO | |
| NUDT8 | chr11 | 67629291 | 67629391 | - | 0 | 0 | 0 | NO | |
| ORMDL1 | chr2 | 189783966 | 189784066 | - | 0 | 0 | 0 | NO | |
| PARP10 | chr8 | 143978227 | 143978327 | - | 0 | 0 | 0 | NO | |
| PIF1 | chr15 | 64823953 | 64824053 | - | 0 | 0 | 0 | NO | |
| RPL26 | chr17 | 8384002 | 8384102 | - | 0 | 0 | 0 | NO | |
| TFF2 | chr21 | 42347644 | 42347744 | - | 0 | 0 | 0 | NO | |
| TSPAN2 | chr1 | 115073138 | 115073238 | - | 0 | 0 | 0 | NO | |
| UNC13C | chr15 | 54408988 | 54409088 | + | 0 | 0 | 0 | NO | 1 of 2 TSS |
| USP39 | chr2 | 85628866 | 85628966 | + | 0 | 0 | 0 | NO | |

#### SUPPLEMENTARY FIGURES

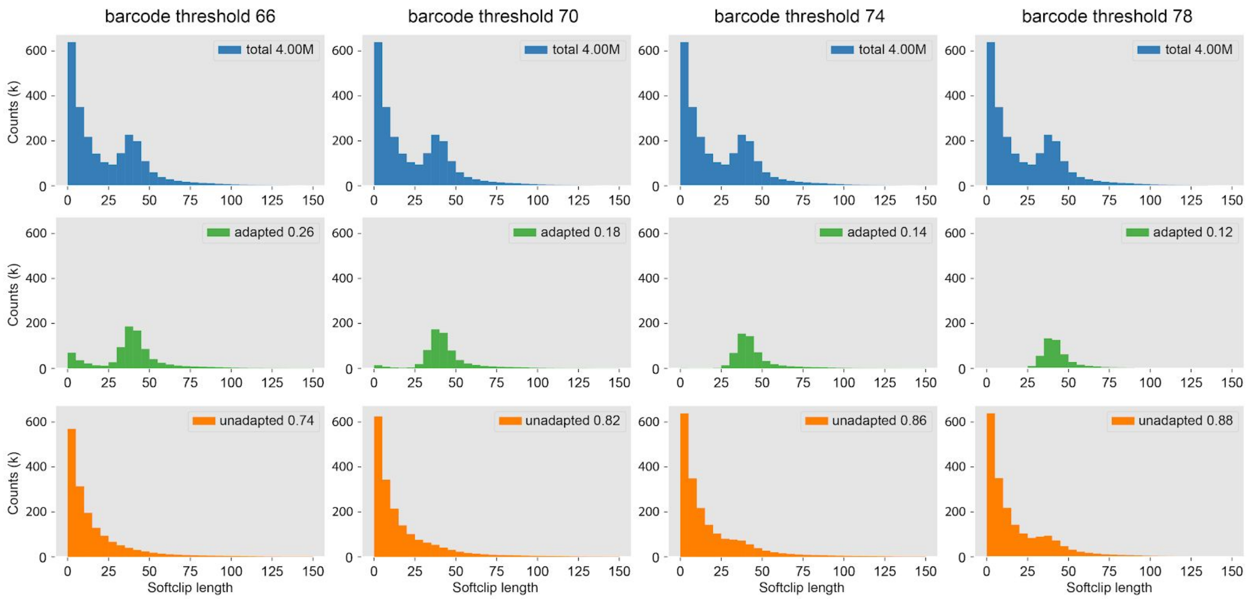


**Supplementary Figure 1** GM12878 Porechop parameter optimization.
The 5' adapter is a 45-nucleotide nt oligomer and is not expected to align anywhere in the human genome, thus it will get softclipped or hardclipped in the alignments. Adapted reads should have a soft or hardclip ~40 nt on the 5' end. Histograms of the 5' softclip or hardclip lengths from untrimmed sequences shows the adapted and unadapted reads. Each column is a Porechop barcode threshold cut off. The top row (blue) are the softclip and hardclip lengths from all the reads. The legend shows the number of reads. The middle row (green) shows the softclip and hardclip lengths from reads identified by Porechop as adapted. The legend is the proportion of cap-adapted reads. The bottom row (orange) are the reads Porechop could not identify the adapter in. We pick a threshold cut off such that the false positives (adapted softclip near 0) and the false negatives (unadapted softclip ~40) are minimized. For GM12878, 74 was the optimal barcode threshold.

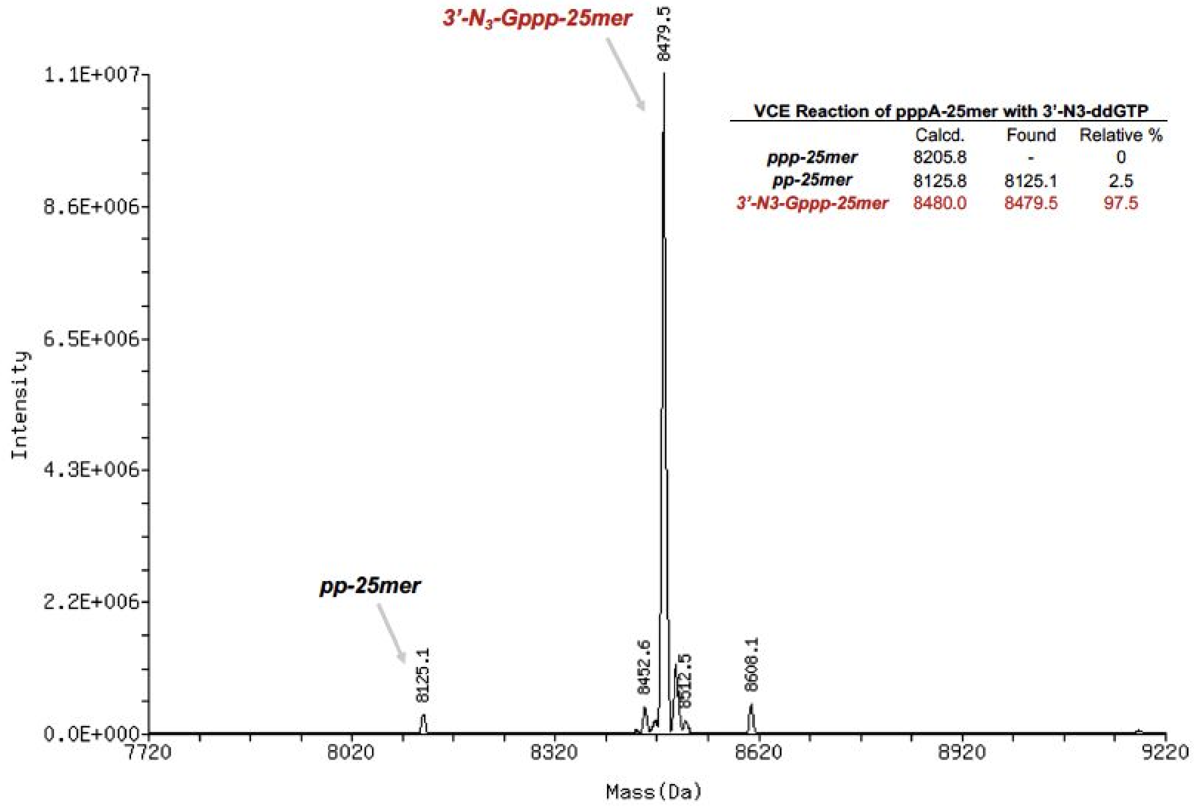


**Supplementary Figure 2** *Vaccinia* Capping Enzyme caps RNA with 3'-azido-ddGTP. Deconvoluted ESI-MS spectra of a synthetic 25-nucleotide 5′-triphosphate RNA oligomer (ppp-25mer) capped with 3'-azido-ddGTP. Tandem Liquid Chromatography-Mass Spectrometry (LC-MS/MS) was performed on a Vanquish Horizon UHPLC System coupled with a Thermo Q-Exactive Plus mass spectrometer operating under negative electrospray ionization mode (–ESI). MS data acquisition was performed in the scan mode. ESI-MS raw data was deconvoluted using Promass HR (Novatia). The composition of each peak was determined by comparison with calculated average atomic mass. The results show nearly complete oligomer capping after 60 min incubation with VCE (see Methods for capping conditions).

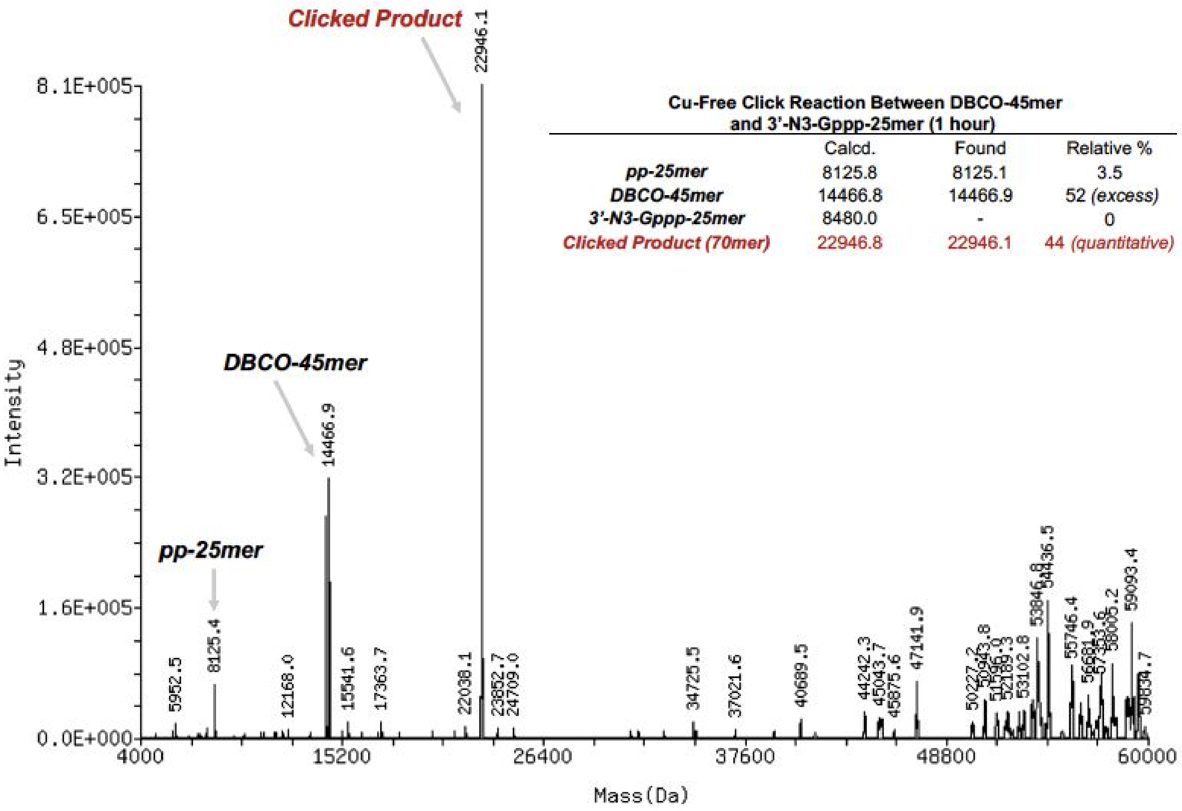


**Supplementary Figure 3** Copper-free Click Chemistry Reaction with a Synthetic RNA Template. Deconvoluted ESI-MS spectra of the synthetic 25-nucleotide azido-ddGTP capped RNA oligomer from Supplementary Figure 4 coupled with the 3′-DBCO RNA adapter (DBCO-45mer). LC-MS/MS and spectral deconvolution were performed as described in Supplementary Figure 4. The composition of each peak was determined by comparison with calculated average atomic mass. The results show that after 60 min the azido-ddGTP capped RNA is entirely consumed forming the desired adapted RNA ("clicked" product). Excess of unreacted 3′-DBCO adapter and some remaining 5'-diphosphate RNA (pp-25mer) were also detected.

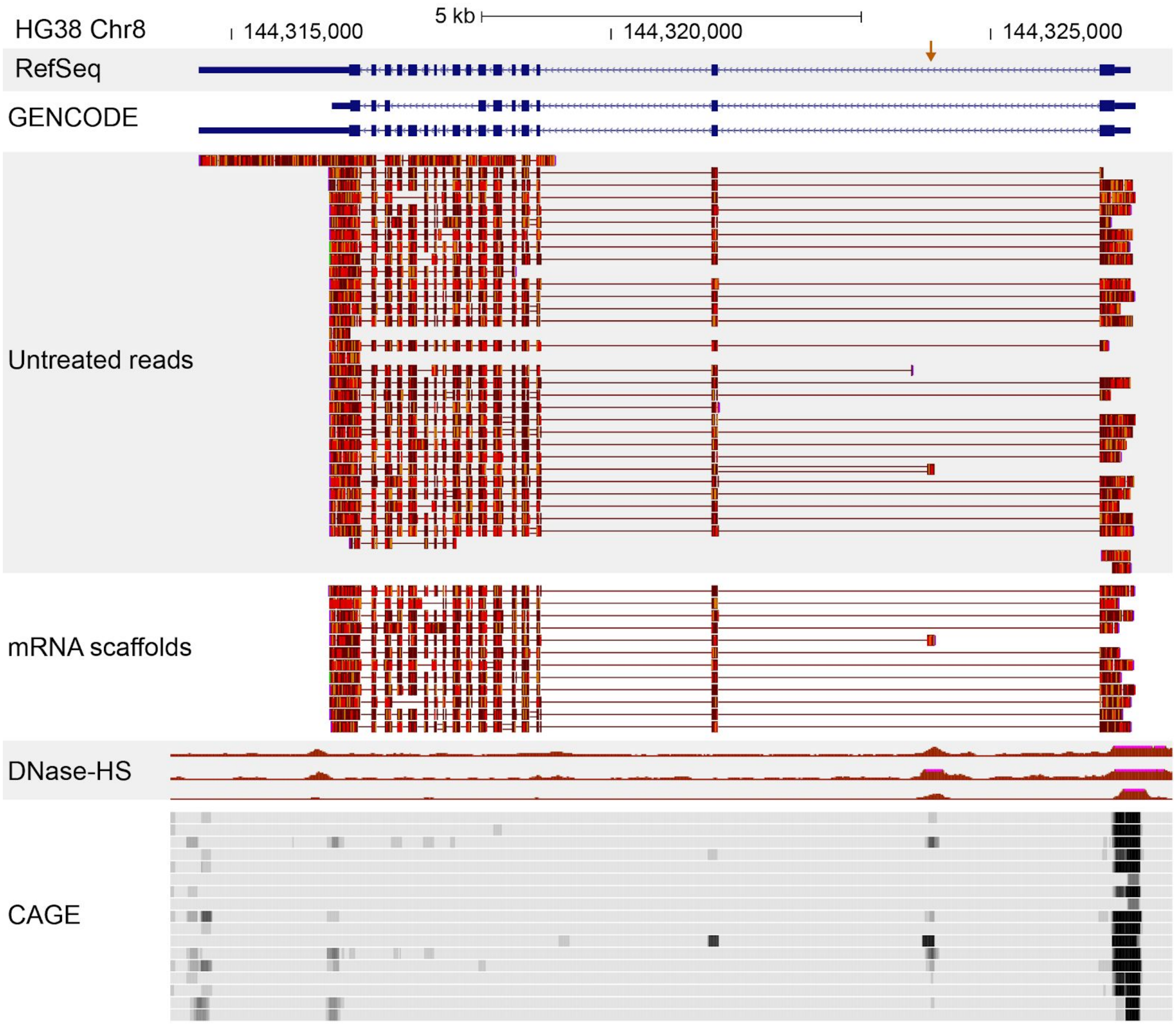


**Supplementary Figure 4** Evidence for an unannotated Diacylglycerol O-Acyltransferase 1 (DGAT1) isoform is supported by a single high-confidence mRNA scaffold. The row entitled mRNA scaffolds includes 12 aligned reads in the 3′-to-5′ orientation. Most of these aligned to a GENCODE v.32 annotated isoform. One of these mRNA scaffolds (orange arrow) corresponds to an unannotated first exon of a proposed unannotated DGAT1 isoform. This unannotated isoform is also observed among the untreated reads. However, untreated reads lack strong evidence of a mature mRNA 5′ end because they are not cap-adapted. The first exon of the proposed unannotated isoform is consistent with open chromatin revealed by the DNase-HS data.

**SUPPLEMENTARY METHODS**

**Synthesis of 3′-Azido RNA adapter**

The 45-nucleotide 3′-azido RNA oligomer (CUCUUCCGAUCUACACUCUUUCCCUACACGACGCUCUUCCGAUCU) was synthesized on an ABI 394 DNA synthesizer (Applied Biosystems) starting with 3'-alkyne modifier Serinol CPG (BaseClick, #BCA-02) and UltraFast RNA TBDMS RNA amidites (Glen Research: Bz-A-CE #10-3003, Ac-C #10-3015, Ac-G-CE #10-3025, and U-CE #10-3030). The oligonucleotide was deprotected according to the manufacturer's protocol using ammonium hydroxide/methylamine and purified using a Glen-Pak RNA purification cartridge (Glen Research, #60-6100) followed by PAGE. The oligonucleotide was further purified by PAGE followed by a desalting step on RP-HPLC (C-8 Higgins Analytical) using 0.1 M TEAB and acetonitrile as the mobile phase. The purified oligonucleotide was converted to 3'-azido in a total volume of 889.2 µL, containing 25% v/v DMSO in 0.2 M triethylammonium acetate buffer, pH 7 as follows (unless other specified, final concentrations are given): 100 µM oligomer, 20 mM $N_3$-PEG1-$N_3$ (BroadPharm, #BP-20908) and 500 µM ascorbic acid were combined and the solution briefly degassed with nitrogen. 44.4 µL of a 10 mM Copper(II)-TBTA complex in 55% aq. DMSO (500 µM final concentration) (Lumiprobes, #21050) was added and the solution briefly degassed with nitrogen. The reaction stirred for 3 h at room temperature in absence of light. The reaction was then dissolved in 0.1 M TEAB (up to 35 mL) and purified by C8 HPLC (Higgins Analytical) using 0.1 M TEAB and acetonitrile as the mobile phase to yield the 3'-azido RNA adapter.

**Adaptation of propargyl capped RNA via Copper-catalyzed Click Chemistry**

Copper-catalyzed click chemistry reactions were performed in a total volume of 10 µL, containing 25% v/v DMSO in 0.2 M triethylammonium acetate buffer, pH 7 as follows (unless other specified, final concentrations are given): 0.5 µM propargyl capped RNA, 4 µM 3′-azido RNA adapter and 500 µM ascorbic acid were combined and the solution briefly degassed with nitrogen. 0.5 µL of a 10 mM Copper(II)-TBTA complex in 55% aq. DMSO (500 µM final concentration) (Lumiprobes, #21050) was added and the solution briefly degassed with nitrogen. The reaction shaken overnight at room temperature in absence of light. The adapted RNA was recovered using RNA Clean & Concentrator (Zymo Research, #R1013).

*Porechop adapter identification* The search sequence TCCCTACACGACGCTCTTCCGA was added to Porechop's adapters.py file as a new 5' barcode. The native RNA nanopore reads had the U's in the sequence replaced with T's using a python script, and then quality filtered using NanoFilt to q7. Porechop was used on the quality filtered reads with the following conditions: --barcode_diff 1 --barcode_threshold 70 (S. cerevisiae) 74 (GM12878) -i file.fastq -b outputdirectory. --untrimmed was used while optimizing the threshold parameter.
